# High-throughput task to study memory recall during spatial navigation in rodents

**DOI:** 10.1101/534313

**Authors:** Lucia Morales, David P. Tomàs, Josep Dalmau, Jaime de la Rocha, Pablo E. Jercog

**Author notes:** These authors contributed equally to this work (Lucia Morales, David P. Tomàs).

## Abstract

Spatial navigation is the most frequently used behavioral paradigm to study hippocampal dependent memory formation in rodents. However, commonly used tasks can present some limitations: i) they are labor intensive, preventing the implementation of parallel testing for high-throughput experimentation; ii) yield a low number of repeated trials, curtailing the statistical power; iii) are hard to combine with neural recordings, because tethering sometimes interferes with behavior; iv) are not based on overt behavioral responses that can be precisely timed, making difficult the identification of the underlying neural events; v) produce a low spatial coverage, limiting the characterization of neuronal patterns related to spatial information. To circumvent these limitations, we developed a spatial memory task that required minimal human intervention, allowed simultaneous and unsupervised testing of several mice, and yielded a high number of recall trials per session (up to ~20). Moreover, because recall sessions could be repeated over many days, the task provided enough statistical power to characterize in detail the animals behavior during memory recall, even to quantify the decay in spatial accuracy of memories as they are stacked across days. In addition, the task is compatible with neural activity recordings. Together, these features make our task a valuable tool to start dissecting the neural circuit dynamics underlying spatial memory recall.

## Introduction

Navigation is of extreme importance for the behavior of many species, and the memory of locations and paths are in some cases crucial for survival. After the discovery of place-cells located in hippocampus (O’Keefe J, Dostrovsky 1971), it was proposed that these cells are the main neural correlate of spatial information. The hippocampus has also been identified for many decades as a key component for the formation of a map of memories, also called “cognitive map” (O’Keefe and Nadal 1978). Experimental brain lesions and inactivations have shown that the hippocampus (Winocour et at., 2001) and entorhinal cortex (Oswald et al., 2003) are the two main structures necessary for spatial navigation, also called “allocentric”. This type of navigation depends on distal cues while egocentric navigation depends on proximal cues. To study how the rodent’s brain creates maps of space, as a model to understand how humans create cognitive maps of memories, researchers have been using spatial navigation tasks to assess behaviorally as well as electrophysiologically the mechanisms behind these processes.

The most widely used tasks to study spatial navigation memories are the Morris water maze (MWM) (Morris 1981) and the radial-arm maze (RAM) (Olton and Samuelson 1976). The RAM task tests long-term memory by counting as errors visits to unrewarded arms, learned on a previous session. To do the task correctly it is also important to avoid re-visiting an unrewarded arm that was previously explored. One disadvantage of using the RAM test is that, once the animal enters an arm, an odor trace creates a local cue that makes it impossible to assess whether the high score is due to actual memory or to a strategy based on avoiding repeating an arm that has been marked by an odor trace. The MWM task, where the animals need to swim in a pool in search of a hidden platform, aims to remove local cues (e.g., odor traces) and force the animal to exclusively make use of distal cues. As a downside, MWM is limited in both: the number of trials that rodents can make per memory test session, and the possibility of the simultaneous recording neural activity. Even that other alternatives (Barnes et al 1979, Post et al. 2011) or modifications of these tasks (Bimonte et al., 2000, Fouquet et al., 2013) have been used with high success in the field (for review see Sharma et al., 2010, Vorhees & Williams 2014), but some of the limitations are maintained. Another widely used task to test spatial memory is the spatial object recognition (SOR) task (Ballarini et al., 2009). In this task the animal is presented with two familiar objects, one of which has been displaced. The memory readout is the time spent with the displaced object, relative to the fixed one. As this follows from natural behavior the animals do not need to be trained, but the large behavioral variability during the time window to measure memory, makes the statistical power of the task weak.

As a consequence of the tasks above not being designed for stereotyped, high-throughput experimentation, establishing correlates between behavior and neural activity is problematic. Previous attempts to correlate memory with neuronal activity (Kentros et al 2004) used recordings from hippocampal neurons while animals performed a spatial navigation task that besides counting with few trials per session, the resulting outcome did not enable measuring the distance to the correct answer. A more complex spatial navigation task was used by Dupret et al., 2010, where animals needed to learn the location of 3 rewarding wells in a lattice of wells. Animals learned to navigate from their “home” station to the rewarded wells over 40 trials per session, and the memory accuracy of the rewarded locations was tested on memory recall session, at 2 and 24hrs. The task allowed researchers to collect neural activity while animals performed the learning and memory recall sessions. This behavioral paradigm despite of having many benefits has two main limitations: (1) labor intensive human intervention, which prevents the use of large animal’s cohorts per day; and (2) the scarce spatial coverage during the task, hampers a good characterization of neural activity (e.g. place-fields) related to the behavior.

Here we introduce a novel behavioral task that is completely computer controlled, where animals perform spatial navigation with a recall session with many trials, that allows to investigate the recall of several different memories learned on different prior days. The behavioral task also allows to record neural activity, without diminishing the performance when animals are connected to the recording equipment.

## Results

### The task

The main goal of the memory task presented here was to obtain high number of trials during a spatial memory recall session with minimal human intervention. The task consisted in a spatial navigation search within a circular arena, to learn and then recall, the location of a rewarding water-spout (port) (Fig. 1a). The behavioral box had transparent walls forming an hexadecagon, to allow the visualization of distal cues. Located on the walls, there were 8 equidistant ports that measured nose pokes and could also deliver water. The control of the ports, as well as the real-time video tracking of the animal was monitored by a computer running an open-source software developed by our group. For this reason, the task did not require any human intervention besides bringing the animals in and out, and starting the behavior acquisition program. The behavioral box was inserted in a sound isolation chamber, which shielded the animals from sound arriving from the room and from neighbor boxes. The chamber also provided distal visual cues, in the form of cards, located in the inner walls (1 cubic meter). Moreover, the isolation boxes allowed us to have several behavioral setups in the same room (see Supplementary Fig. 1). In total, our current configuration allowed a single experimenter to run three animals in parallel simultaneously, summing up to 18 animals on a ~7 hours working day.

**Figure 1:**
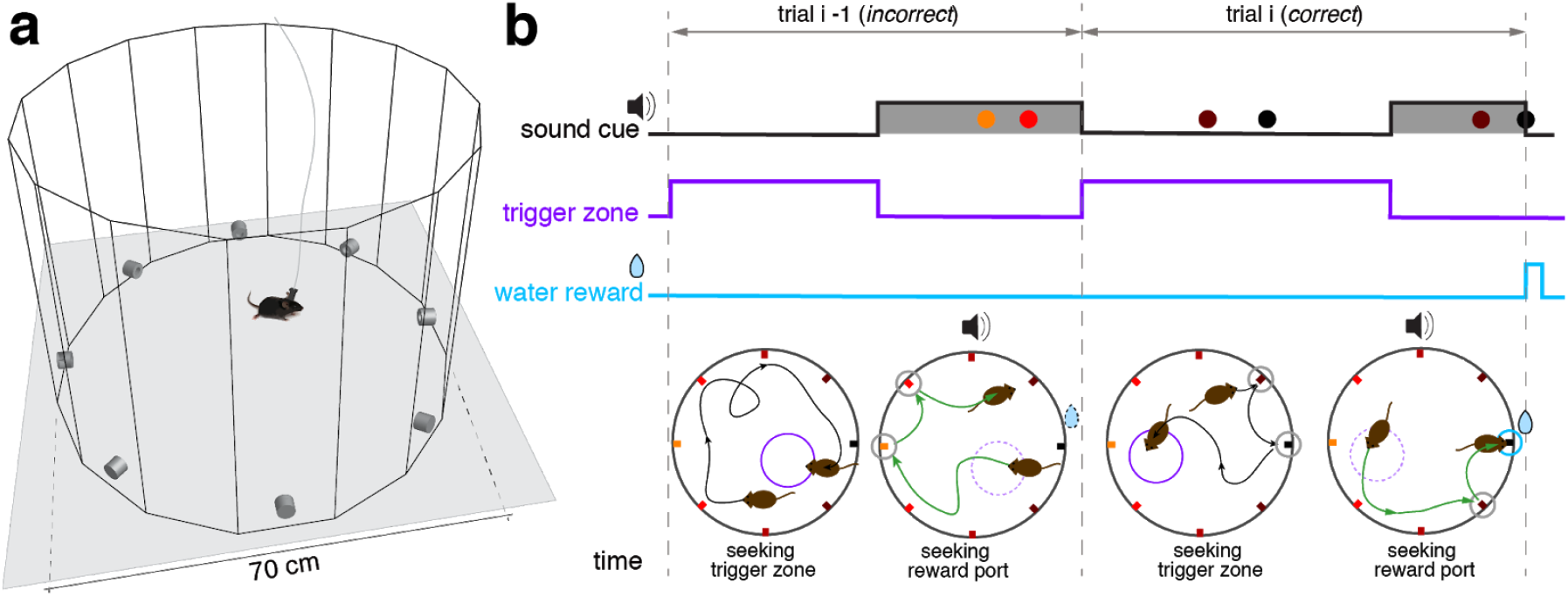
Trial structure of the spatial memory task. **a)** Schematic of the open field built in transparent acrylic with 8 water spouts (ports) controlled by a Python-based software coupled to Arduino boards. The arena is placed inside a 1 cubic meter sound isolation box with cards placed on the walls serving as visual distal cues for the animals. **b)** Schematic of two consecutive trials during a learning session of the task. Top traces show the temporal structure of the sound, trigger zone activation and reward, along with the pokes in the different ports (colored dots) during two consecutive trials. Bottom diagrams show the trajectory of the animals during the two trials and the location of the ports in the maze. Ports are colored according to distance to the correct port (black port at NE). In the first part of each trial, animals walk around the box seeking to start the reward window during which water was available (black trajectory in bottom panels). Reward window starts when they step into an invisible trigger zone, randomly placed in a different location on every trial (small violet circle in bottom diagrams) and it lasts up to 6 seconds. During this window a sound is played cueing the mice to try to reach the correct port (green trajectory). If the animal does not reach the correct port during this window, perhaps because it pokes in other non-rewarding ports, the trial is considered incorrect (trial *i* −1 in b). In contrast, the animal reaches the correct port, independently of whether it poked in incorrect ports before (typically close to the correct port), it receives the reward (10 μ L of water), the sound stops and the trial is considered correct (trial *i* in b). The correct port is fixed in each day (learning and recall sessions), but changes randomly from day to day.

Every day, the task was composed of two sessions: a learning and a recall session. During the learning session, lasting 15 mins, animals learned which of the ports gave reward that day. During the recall session (10 mins long), which starts 2 hours after the training session, we tested the memory of the rewarding port location by removing the reward during a variable time interval at the beginning of the memory session. Except for the absence of reward, the trial structure of the two session types was identical.

### Learning Session

The learning session started after each animal spent ~100 seconds of acclimatization exploring the arena. To initiate the experiment animals seeked to activate the reward availability by stepping on an invisible trigger zone that was a disc of a fraction of the arena’s area (1/16), which was randomly placed in different positions on each trial (Fig. 1b). When the animal walked into the trigger zone, the reward window started lasting up to 6 seconds and a sound was played cueing the animal about the availability of reward at the correct port. The cue was a pure tone above 8kHz (different freq. at different behavioral setups) played at 40 dB. A trial was considered correct if the animal poked into the correct port during the reward window, even if he poked in other ports prior to the correct (see e.g. *trial-i* in Fig. 1b). The reward window was interrupted when the water reward was harvested (see the correct poke closing the reward window in *trial i* in Fig. 1b). Error pokes were never punished. Trials in which the animal did not poke into the correct port during the reward window, were computed as incorrect trials (see *trial i-1* in Fig. 1b).

We leveraged on the task feature that required animals to make overt and discrete responses to obtain reward and used the spatial distribution and timing of poke counts to characterize the behavior during the learning sessions. Poking patterns in each session could be visualized by drawing poke raster plots (Fig. 2a). The mice goal was to identify the rewarding port as fast as possible. To achieve this, they started by poking at the maximum number of ports during the sound cue (Fig. 2a bottom of raster plot; see also Example 1 in Fig. 2b). Once they obtained reward in a few trials (e.g. 2-3), they were able to generate trajectories with almost no errors (Example 2 in Fig. 2b; Fig. 2d left). When they made errors, these were typically on the first neighboring ports relative to the correct one (Examples 3 and 4 in Fig. 2b). To summarize the poking statistics during the entire session, we built poke histograms (Fig. 2d right). The narrower the poke histogram around the correct port was, the more spatially precise was the animal at solving the task. This allowed for a fine quantification of the accuracy of the animal’s navigation and memory, a feature we will exploit in the quantification of the memory accuracy during recall sessions. Because animals had to explore the maze in order to find the trigger zone, the task provided a dense spatial coverage of the arena, a requisite for a good characterization of neural activity encoding space features (e.g. place cells or grid cells). Animals tended to spend slightly more time along the maze perimeter close to the ports, especially near the rewarding port, but they also covered the open space uniformly (Fig. 2e). This uniform coverage of the maze was induced by the uniform distribution of trigger zones, that forced the animals to visit the whole surface of the arena (Fig. 2f).

**Figure 2:**
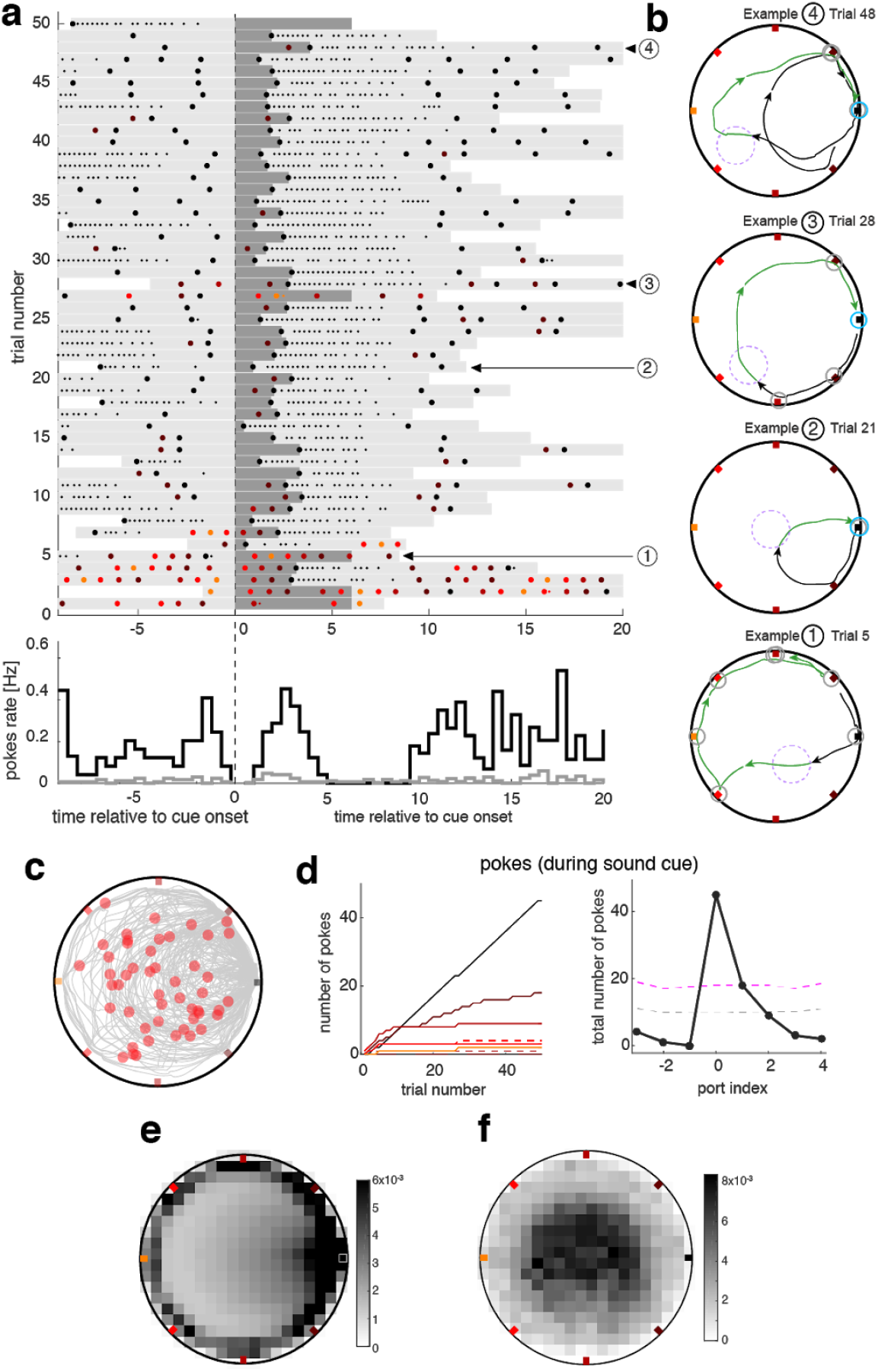
Mice behavior during the learning session. **a)** Top: pokes raster plot during an example learning session shows the timing of pokes at different ports (colored dots matching the port color code) ordered in time by the sound cue onset for each trial (vertical axis). Dark gray bars represent the sound cue duration. Light gray bars indicate the duration of the trigger zone seeking period of the trial. Small dots indicate persistent licking in a given port. In this session the animal seeks the correct port during the first five trials after which he starts to accurately find it in almost all consecutive trials. **Bottom:** pokes cue-triggered time histograms for correct port (black line) and the average over all incorrect ports (grey line). The time bin for the poke rate is 0.5 secs. **b)** Trajectories during four example trials. Color code of pokes and trajectory parts as in Fig. 1b. Trials at the beginning of the session were more explorative (Example 1) until the animal unequivocally identified the correct port location (e.g. in trials 6-9 in this session). From that point on, trajectories were either directed towards the correct port (Example 2), or with only one error poke during the sound cue (Examples 3 and 4). **c)** Trajectory of the animal during the entire session. Red dots represent the position of the animal at sound cue onset. **d) Left:** cumulative poke count vs. trial number for each individual port during sound cue. Correct port (black line) shows higher poke count than any other port during tone. **Right:** poke histogram during sound cue, for each port index ordered by the distance to correct port. Magenta dashed line represents the significance level (*P* < 0.01) over which poking probability was significantly larger than that expected from a uniform distribution Gray dashed line is the mean value of the uniformly shuffled data. e) Density map of the time spent on each spatial bin, for all animals and all sessions oriented with the correct port on the East direction. Density is higher around the correct port, as well as on the other ports, but the coverage inside the arena is quite dense. f) Density plot of distribution of positions where trigger zone is intersected by the animal trajectory, for all sessions and all animals.

Several aspects of the behavioral analysis suggested that animals were navigating in the arena in order to solve the task. Average performance, defined as the percentage of correct trials, rapidly increased with the number of trials until it reached a plateau around 85% (Fig. 3a). In general, animals reached 80% performance in less than 20 trials. The animals became faster in finding the correct port during the session (Fig. 3b) obtaining the reward in approximately 2.5 seconds. As the session progressed, they also made fewer errors during the sound cue, indicating that their spatial accuracy increased with trial number (Fig. 3c). Interestingly, the average time to find the trigger zone also decreased significantly along the session, possibly due to an increase in task engagement as the session progressed. All together this shows that animals were able to excel the task within the early part of each session.

**Figure 3:**
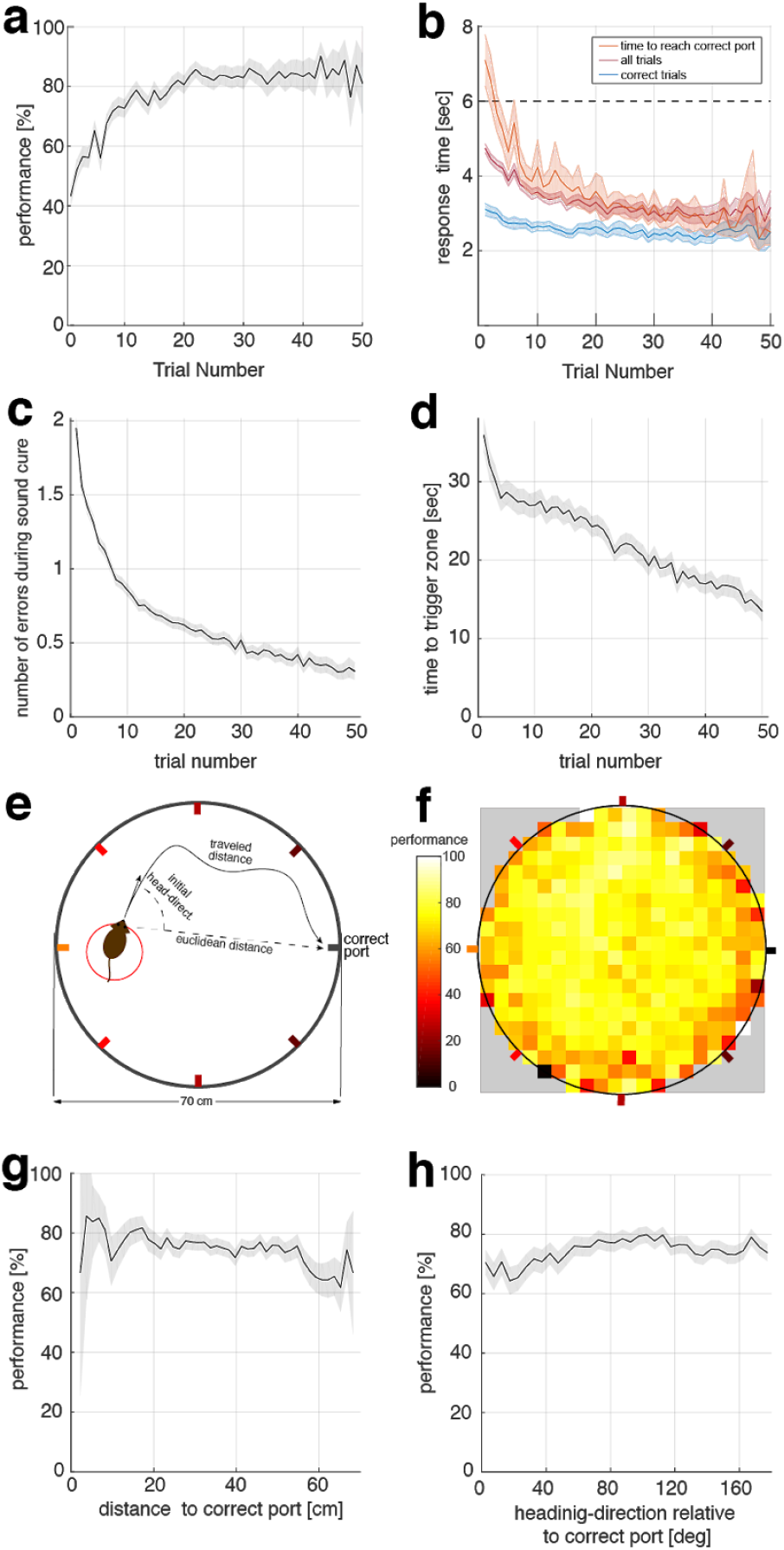
Spatial navigation is used to solve the task. **a)** Performance versus trial number, averaged for all animals (n=21 mice) and sessions (black line). Shaded areas represent the s.e.m. **b)** Time to find correct port in three different conditions: for correct trials only (blue), for correct and incorrect trials (magenta) (incorrect trials response time = 6secs), and total time to reach correct port (orange) even after the 6secs sound cue window. The time to find the correct port decreases with the trial number. **c)** Number of errors during sound cue, is the number of ports poked before poking correct port. Errors decreases with trial numbers. **d)** Time to find trigger-zone. After sound cue is finished, we computed the time from this moment until the animal hits the randomly placed trigger zone. Time to find trigger-zone decreases over time. **e)** Schematic of the navigation to describe the initial heading direction angle and initial euclidean distance to the correct port when the animal hits the trigger zone. **f)** Map of performance outcome for a given trial, depending on the initial position on the arena. Performance does not change depending on the initial position in the arena. **g)** Performance is independent of the initial euclidean distance from the animal to the correct port. **h)** Performance is independent of the initial heading-direction angle relative to the correct port. Error bands in all panels indicate s.e.m.

One strategy that animals could employ to solve the task could be the use of stereotypical trajectories repeatedly traced on the arena over multiple trials (i.e. trails). Such a strategy should yield strong dependency of the performance on the initial position of the animal relative to the port. We found however that performance did not depend substantially on the initiation point of the trajectory from the trigger zone towards the rewarding port (Fig. 3f). We also assessed whether performance varied according to the distance from the initiation point to the correct port or on the heading-direction angle relative to the correct port at the initiation point (Fig. 3e). We did not find substantial differences in performance related to either of these factors (Fig. 3g and h). Together, these analyses suggest that animals did not use a stereotypical strategy to solve the task, but that their strategy was flexible enough to find the rewarded port in a diversity of conditions imposed by the randomization of trigger zone location.

### Recall Session

Two hours after the learning session, animals underwent a recall session to measure if they remembered the correct port location. During the first part of the recall session animals did not receive reward if they poked in the correct port. The sound cue did not stop when they poked in the correct port either. The absence of feedback was aimed to prevent animals from re-learning the position of the correct port. To maintain the motivation of animals to seek reward during the memory recall session, reward was omitted only in a fraction of sessions and only during a variable time which could be 1, 3 or 5 minutes (set randomly). When the animal lost the motivation on a given recall session, water was made available on the following days during the entire recall session until motivation was recovered. In these conditions of both spatial and temporal uncertainty, mice seeked for the reward during several trials (e.g. 10-20) during which they persisted in trying to find the rewarding port. These trials were used as equivalent repeated memory probes. We measured the recall of the spatial memory by quantifying the probability to poke in the different ports during trials in which water was not available (e.g. trials 1-16 in the example shown in Fig. 4a). Despite the large variability in their poking behavior, animals exhibited a weak but significant tendency to preferentially poke the correct port learned 2 hours before (Fig. 4b-e). Once water was introduced and animals obtained reward in one trial, then they focused on almost exclusively poking the rewarded port without much behavioral variability (Fig. 4a-b, d-e, blue rectangle) as they did in the last trials of the learning sessions.

**Figure 4:**
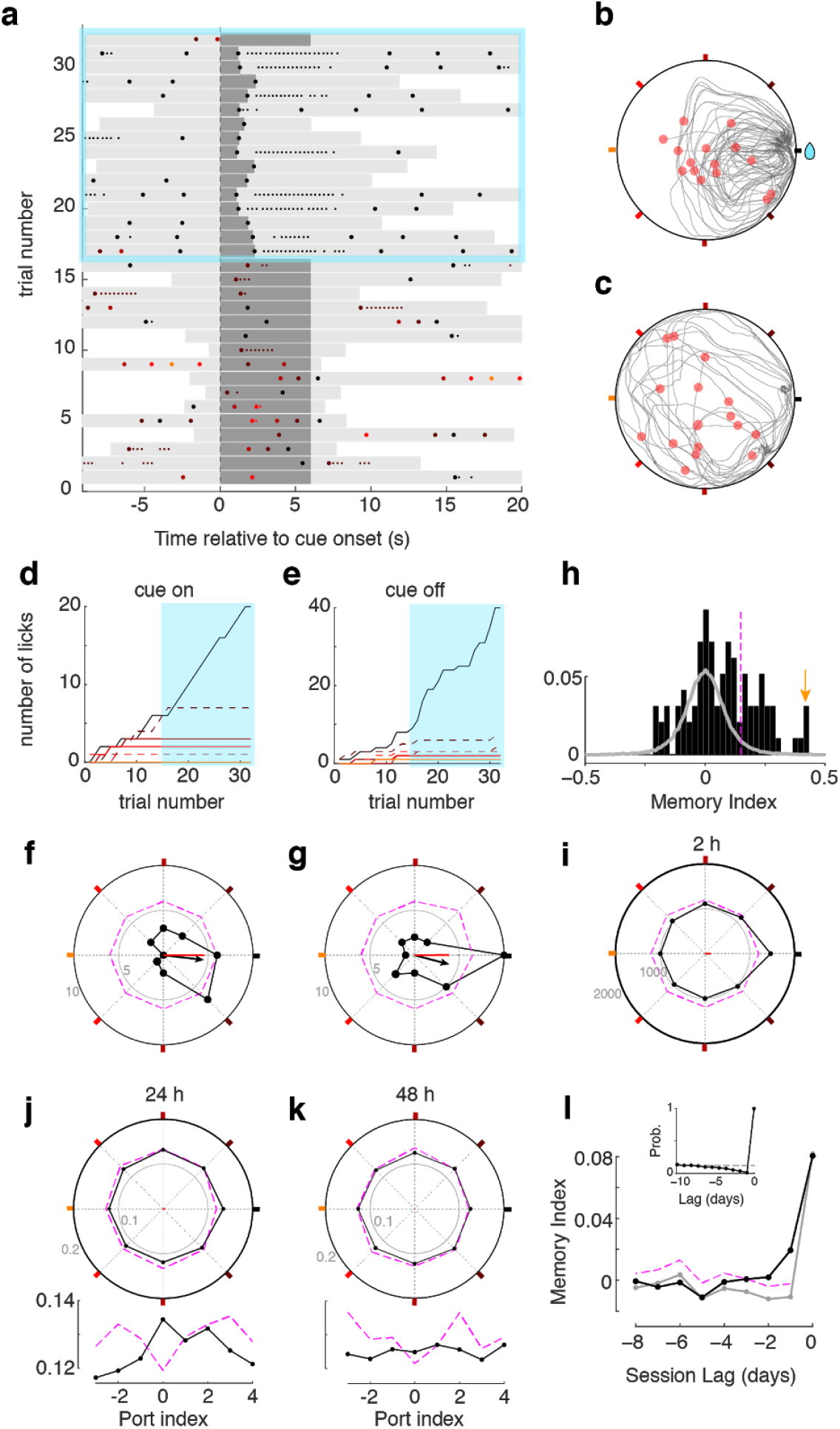
Mice behavior during memory recall sessions. **a)** Pokes raster plot during an example memory recall session performed by mouse 4032, displayed as explained in Fig. 2a. During recall sessions, correct pokes during the first *n* minutes of the session (*n* = 0, 1, 3 or 5) were not rewarded, to avoid relearning of the reward location. Sound cues were never interrupted during this period. After the *n* minutes, correct pokes were rewarded as during the learning session (blue rectangle). In this example session, during the first *n =* 5 minutes, corresponding to trials 1-16, water was not available, whereas it was in trials 17-31. **b-c)** Trajectory of the animal during the same session for the period without water (c) and water (b). Representation is equivalent to that shown in Fig. 2b. **d-e)** Cumulative poke count versus trial number for each port during sound cue (d) and off the sound cue (e). Correct port (black) and one neighbor port (dashed brown) show higher poke count in both cue on and cue off condition during the period without water. Once water was available and harvested (blue area), only the rewarded port was poked. **f-g)** Poke histograms in polar coordinates for the example session (black dots), showing the number of pokes in each port as the radial distance of each point to the center, for the on cue (f) and off cue (g) conditions. Colored squares mark the angle of each port (as in Fig. 1b). Small gray numbers show the count number of the radius of the inner and outer circles. Dashed magenta line shows pointwise significance bound obtained from shuffled data (*P* < 0.01, one-tailed; see Methods). Black arrows show the vector summation of the counts over all the ports. Red lines show the MI for the session. **h)** Histogram of MIs obtained from individual recall sessions of mouse 4032 (black bars). Gray histogram shows the MI values from the surrogate data set drawn from a uniform poking distribution. Magenta vertical line shows the significance bound of individual session MI (*P* < 0.01). Yellow arrow marks the session shown in panels a-g. **i)** Accumulated poke histogram for all sessions (n = 97) of mouse 4032 considering both on and off cue conditions. Magenta dashed line indicate significance bound (*P* < 0.01, one-tailed). Red line shows the MI over all sessions. **j-k)** Normalized poke histograms aligned at the port learned 24 hours (j) and 48 hours (k) before (black dots) show a significantly increased preference to poke in the ports memorized in the two previous days (*P* < 0.01, permutation test). Histograms were the average across animals (n= 19 mice) and sessions (mean no. sessions 40; range 97-8). Lower insets show the same poke histograms unfolded for finer visualization. l) Averaged memory index MI versus the session lag (black dots, n= 19 mice) shows that there was significant memory recall of the correct ports from the three previous sessions (i.e. up to 72 hours). Significance was assessed generating a surrogate data set with *only* 2h memory that followed the same sequence of rewarded ports across days (mean across surrogates is shown in gray). Confidence bound for lags ≥ 24 hours is shown in magenta (*P* < 0.01, one-tailed). *Inset:* probability that the correct port from a given day was also selected as the correct port in previous days.

### Analysis of memory recall accuracy

We developed a *memory index* (MI) to quantify the recall accuracy in single sessions, or averaged across sessions. To compute the MI, we first created a spatial poke histogram represented in polar coordinates. This choice of coordinates easily illustrated the tendency of the animal to poke the ports in each orientation of the box perimeter (e.g. plots Fig. 4f-g show a tendency to poke in the East and SE ports; correct port at East). We then built a pokes vector by performing a vector sum with the contributions of each of the eight ports, each pointing in a different direction, and normalized by the total number of pokes (see Methods). The resulting vector summarized the excess of probability to poke in a certain direction (see black arrow in Fig. 4f). By projecting the vector onto the x-axes we obtained the MI which quantified the net poking preference towards or away the correct port (red line in panels Fig. 4f-g). The MI ranged between +1 (all pokes were in correct port) and −1 (all pokes were in the port opposite to the correct port). The MI was therefore based on the poking probability in each of the ports and not exclusively in the correct, what conferred to the MI a robust statistical power. Moreover, the MI was independent of the total number of pokes during the session, allowing the comparison across sessions and animals, typically yielding a different number of pokes.

Figure 4f-g shows the poke histogram and the MI of an example mouse in a session with a particularly high recall accuracy. Across sessions, this animal exhibited large variability in the MI, with 31 out of 97 recall sessions yielding a MI significantly different from zero (*P* < 0.01; see black histogram in Fig. 4h). To test for significance, we compared this histogram with the distribution of MIs obtained from a surrogate data set, that lacked any spatial preference: using the same number of pokes per session than the original data. The surrogate data were generated following a uniform poking distribution across ports (gray histogram; vertical dashed magenta line shows the significant level at *P* < 0.01, one-tailed; see Methods for details). Capitalizing on the design of the task, memory accuracy could also be quantified by pooling sessions performed at different days with different correct ports. For this, we pooled the poke histograms from multiple individual sessions separately (Fig. 4i shows animal 4032). Statistical significance was then assessed by shuffling the values of each individual session histogram and then summing across sessions to obtain a pooled shuffled histogram (dashed magenta line shows *P* < 0.01 level). Pooled poke histograms for each of the mice tested are shown in Supplementary Fig. 2. From the pooled histograms, a MI was obtained for each animal, as done from individual session histograms: 15 out of 19 mice showed a significant memory index (*P* < 0.01, one-tailed permutation test). In sum, our metric of recall accuracy, the memory index MI, allowed us to show that the majority of our animals were able to show significant recall when averaging across sessions.

### Assessing memory recall from previous days

Despite the ability of mice to remember the port that provided reward during the learning session 2 hours before, their poking behavior was still quite variable. We thus asked whether this variability could be caused, at least in part, by the recall of the memories acquired in previous days. For instance, in the example recall session shown in Fig. 4f-g, besides the increased frequency to poke the correct port (East position), the animal also showed a strong tendency to poke the SE port (Fig. 4f-g). It turned out that this port was the correct port during the previous day. Was this deviation towards ports learned on previous days systematic across animals and sessions? Our task was well suited for this question because animals underwent dozens of learning sessions over several months, so that the learning of the procedure was well established and sessions could therefore be taken as a series of concatenated learning-recall experiments. For each animal, we built a poke histograms by pooling all sessions, but aligning each session to the correct port learned *n* days ago (*n* = 1, 2, …8). We then normalized these histograms and took the average across animals to maximize the statistical power to detect significant deviations in these histograms (Fig. 4j-k) and in the corresponding MI (Fig. 4l). To assess the statistical significance of the MI from previous sessions, we built a surrogate data set composed of the same number of sessions and pokes, that yielded the same 2 hours MI than the original data but in which, by construction, there was not any trace of previous memories (Fig. 4l gray line; see Methods). From this surrogate data set, we obtained the significance bound of the MI as a function of the session lag (Fig. 4l, magenta dashed line). We found that the animal-averaged MI was significantly greater than the surrogate set for 24 hours (*P* < 0.01), 48 hours (*P* < 0.01) and 72 hours (*P* < 0.05) (compare black and magenta lines in Fig. 4l). For longer session lags, the MI was not statistically different from the surrogates (Fig. 4l). Thus, the large number of recall sessions yielded by our automatized task, with multiple repeated trials per session, allowed us to reveal the memory trace caused by sessions from previous days, up to 72 hours into the past. This finding opened the door to address fundamental questions on how stored memories may interact during the retrieval process (e.g. memory swaps, memory attraction, etc).

### Neuronal activity recordings while animals perform the task

Having shown that our task can quantify the behavioral traces of multiple memories learned over the last few days, we wanted to demonstrate, as a proof of principle, that animals were able to perform the task while their neuronal activity was recorded. We implanted miniaturized-microscopes (nVista) in two animals and recorded the calcium activity from a few hundred neurons in the hippocampal area CA1 during 15 learning and recall sessions (Fig. 5b-d). Performance during sessions in which the two mice carried the mini-microscopes was not different from sessions in which they were untethered (Fig. 5a). We collected the data using an independent computer than the one controlling the behavioral box, and we synchronized off-line the behavioral movies with the calcium frames via the TTL signal from the mini-microscopes stored by our behavioral control program. Analyses performed offline using state-of-the-art methods, we re-aligned frames from the mini-microscope recordings to overcome the deformation suffered by the individual images due to brain movement when the animal ran during the task. The selection of individual neurons boundaries to extract individual calcium activity is performed by an EM algorithm called (CELLMAX) that fits a model of photon emission generated by individual neurons shapes (see Methods). An example picture of the spatial layout of the neurons sorted from the calcium activity movies while the animals performed the task is shown in Fig. 5b. After the preprocessing of the movies, we obtained traces of calcium activity from each individual neuron (Fig. 5c) that were match-up with the animals position and pokes timestamps during the task, to study correlations between neuronal activity and behavior. As an illustration of the quality of the neuronal data, we plotted the response from several neurons during one of the learning sessions (Fig. 5c). Given the dense spatial coverage obtained in the behavioral task (Fig. 2e), we are able to characterize place-cell response patterns with spatial information above 2 bits/sec (Fig. 5d), which is a typical threshold used to study place-cells from electrophysiological recordings methods in freely moving open fields. Overall, this example demonstrates that the behavioral task was suited for the monitoring of large neuronal assemblies using modern calcium imaging techniques allowing for interrogation about the changes undergone by hippocampal circuits during memory formation and recall.

**Figure 5:**
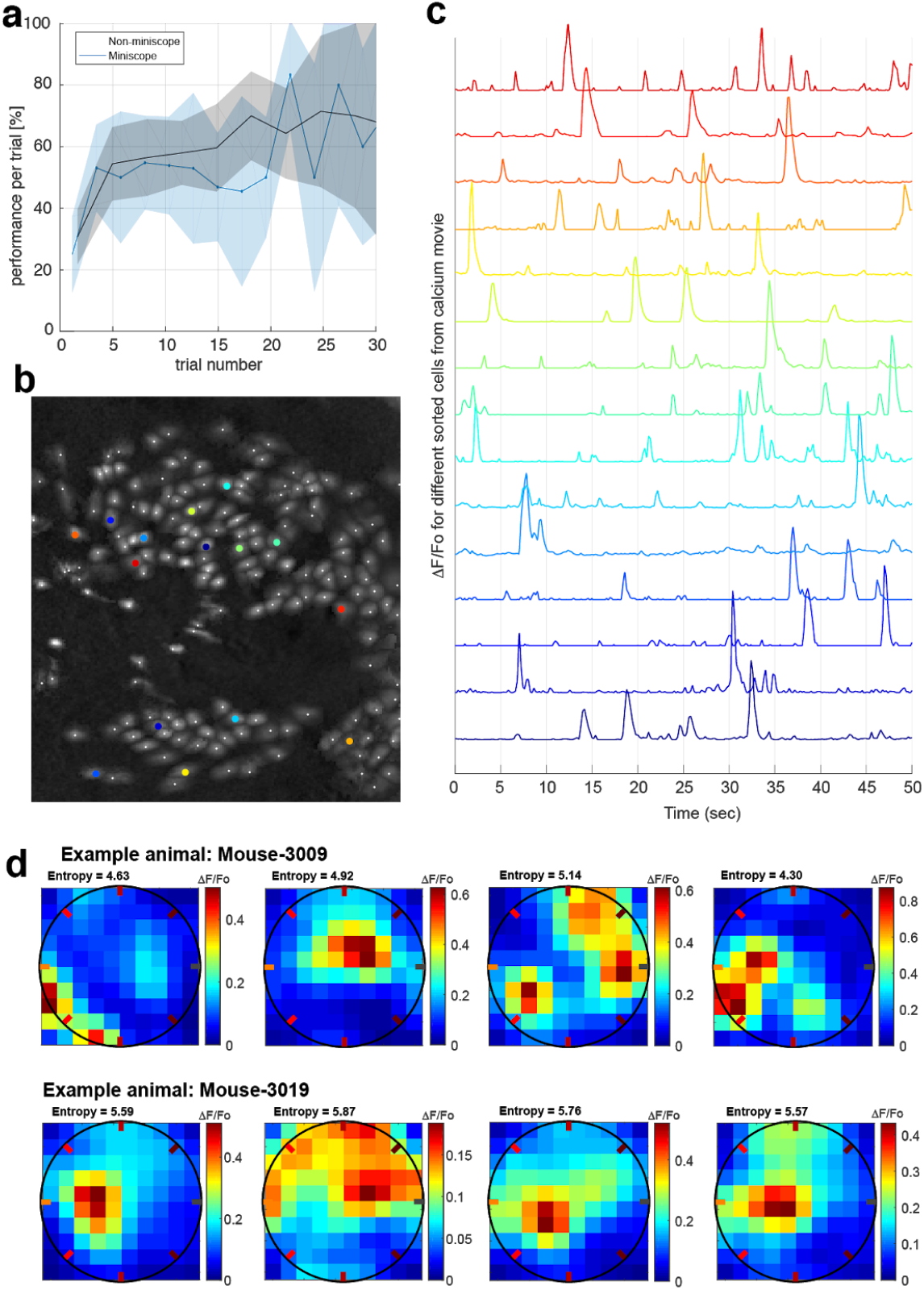
Calcium imaging recordings while animals perform the task. **a)** performance versus trial number does not show a significant difference if we compare the conditions with the animal carrying the mini-microscope (*P* = 0.5, one-tailed t-test). **b)** Example frame calcium imaging registration of the cells recorded simultáneusly while the animal is performing the task (approximately 180 neurons recorded). **c)** calcium activity traces of relative fluorescence (Δ F/F_o_) recorded from selected neurons (colored in b). Traces were processed off-line and used to decode different behavioral variables of the task (e.g. animal’s position). **d)** Example place-fields obtained from two different animals while performing the task. Place-fields were chosen from the distribution of spatial information measured by the entropy of the place-fields neuronal tuning. Note: all recorded place-cells have entropies higher than 2.5 bits/sec.

## Conclusions

We presented a novel memory task characterized by (1) a complete automatization of the task, which did not require almost any human intervention (just for placing and removing the animal from the arena); (2) a trial-based structure in which memory recall was repeatedly probed in each session (up to ~20 recall trials) and animals underwent dozens of recall sessions over the course of several months; (3) overt recall responses in the form of precisely timed nose pokes in the different ports.

The automatisation of the task facilitated a high-throughput behavioral approach (Aoki et al 2017; Han et al 2018), allowing one experimenter to run several experiments in parallel in different boxes (Supplementary Fig.1). Unsupervised behavioral testing reduces the costs of human intervention, decreases the manipulation of animals minimizing stress, and has the advantage of removing the experimenter’s subjective component, both during training and when quantifying behavior. Moreover, behavioral automation requires the standardization of training protocols (e.g. setting well defined criteria about when animals must progress in the training), ultimately increasing the overall reproducibility of the data. Our design should be viewed as an important step towards a fully automated behavioral control and monitorization of the task. Future implementations should consider self-paced sessions, where animals could voluntarily access the arena from their home cages to perform the task when they had the necessity to drink water (Rivalan et al. 2017). Furthermore, memory recalls could be then automatically schedule at different time delays.

Behavioral assays in mice are typically hampered by the large and unexplained variability exhibited by animals during and across repeated experiments. During recall sessions, our mice exhibited a large variability in their search behavior (Fig. 4a): they tended to poke multiple ports in a seemingly random order. By structuring recall sessions in a sequence of repeated trials, we were able to accumulate enough statistics and get around this intrinsic limitation. Because trials were self paced by the animals, motivation across trials is similar and there was no need to make assumptions about when animals stop being engaged in the task. In fact, maintaining the motivation over multiple trials required creating the conditions in which animals were uncertain about the absence of reward during the repeated recall trials. This was done by randomly changing in recall session, the length of the no reward period from 0 to 5 minutes. Had we maintained the duration of this period fixed, mice would have learned that recall sessions did not provide reward (at least for a few minutes), and they would have not performed as many recall trials. Moreover, having the possibility to run the experiment over dozens of concatenated sessions, not only allowed us to achieve a substantial statistical power in our quantification of memory accuracy (Fig. 4), but it opened the possibility to study a repertoire of questions regarding the interactions between memories: can stored memories be swapped during recall? Can the trace of a newly formed memory be affected (e.g. spatially shifted) by the existence of similar memories? Is there a relation between the acquisition of new memories and the forgetting of old ones? Using pharmacological manipulations (e.g. NMDA receptors blockers) or using the task with animal models of inducible amnesia, could further dissect the mechanisms at play during the recall of memories stacked across time.

Finally, our task required animals to learn to poke in the different ports to obtain reward, in contrast to other tasks that make use of natural behaviors which do not need training. Once animals learned to perform the task (which typically took 20-28 training sessions), the behavioral output was delimited to a discrete series of overt responses, i.e. the nose pokes. This feature, not only facilitates the interpretation and analysis of the data, but because pokes are precisely timed, it allows for the synchronization of optogenetics manipulations, inhibiting or exciting different brain areas at precise times around these memory recall events. Moreover, having a well timed behavioral output, facilitates the use of electrophysiological or calcium-imaging recordings aiming to characterize the neural correlates underlying the generation of memory recall event. In sum, this novel memory task has the potential to become an instrumental tool to investigate the processes of memory formation and recall and to dissect the neural circuits dynamics underlying these functions.

## Acknowledgments

We thank Rosa Rodriguez for technical assistance.

LM and DT research was supported by Fundació CELLEX.

JD research was supported by Instituto Carlos III/FEDER (FIS PI14/00203; PIE 16/00014); AGAUR (SGR93); CERCA Programme/ Generalitat de Catalunya, La Caixa Foundation Health Research Award, and Fundació CELLEX.

JdlR research was supported by the European Research Council (ERC) under the European Union’s Horizon 2020 research and innovation program (PRIORS; No 683209) and the Spanish Ministry of Economy and Competitiveness together with the European Regional Development Fund (SAF2015-70324-R).

PEJ research was supported by Fundació CELLEX, and Marie Curie Fellowship FP7-PEOPLE-2013-IIF, N°=627457, (ISOLM).

## Contributions

LM, JdlR, JD and PEJ designed the experiment. LM and PEJ built the behavioral boxes and the isolation chambers. PEJ developed the behavioral control software. LM and PEJ performed the behavioral experiments. PEJ performed the calcium imaging experiments. LM, DT, JdlR, and PEJ analyzed the data. JdlR and PEJ wrote the manuscript. LM, DT and JD helped writing the manuscript. PEJ and JD supervised the project.

## Methods

### Behavioral Box

the arena was a Hexadecagon with 70 cm diameter, made of exchangeable rectangular vertical panels (Fig. 1a), where in every other panel there was a standard water port from Sandworks. LLC, model called “mouse port assembly”. The 8 ports were connected to the computer via an arduino “Mega”, using the serial port. We collected data from the ports at approximately 100 Hz (10 ms interval between samples). A GUI in python displayed all the behavioral readouts in real-time for performance monitoring during learning and recall sessions. During pre-training, the GUI also gave the alarm when the criteria was reached to pass the animals to the next stage (read *pre-training* detailed explanation below).

### Behavioral paradigm - Learning session

All animal procedures were approved and executed in accordance with institutional guidelines (Generalitat de Catalunya: Autorització de projecte d’experimentació N°=9997), and the Comitè Ètic d’experimentació Animal (CEEA) at the University of Barcelona, N^o^=121/18. We used 21 C57BL6/J male mice (Jackson Labs; 9–10 weeks old) that had restricted access to water for a week, drinking twice a day a total amount of approximately 1 mL of water until their body weight is maintained at 80% of the original weight. After reaching the last stage of the pre-training, animals recovered around 95% of their original body weight by only drinking during the two sessions a day. Experiments were ran 6 times per week. Some animals run the experiment for up to 240 days, without showing signs of stress or dehydration.

Animals were introduced in the arena by the experimenter, who also started the program that recorded the behavior’s video, tracked the position of the animal in real time, and collected the nose pokes in each of the 8 ports. The code also recorded the time-stamp of the calcium frames (or the electrophysiology TTL), when recordings were acquired. Mice had 1.6 minutes for acclimatization after they were introduced in the arena. After that, the first trigger zone (a disk of 17.5 cm diameter) was generated and placed randomly (with a uniform distribution in the circle) within the limits of the arena. Animals were trained to find the rewarded locations while the sound cue was active (max. 6 secs). Once the animal found the correct port, they remembered the correct port for the rest of the session (making few mistakes sporadically, see raster Fig2a). Once the animals collected all the reward, they started moving in random trajectories within the enclosure to trigger a new sound cue that signaled when reward was available at the correct port. The sessions lasted for 15 minutes beyond the acclimatization time, and animals made between 30 and 100 trials depending on the animal and the day.

### Behavioral paradigm - Recall session

animals were introduced in the arena by the experimenter. The animal had 1.6 minutes for acclimatization. After that period, the first trigger zone was generated randomly within the limits of the arena as in the learning session. Water was not available during the recall session for the first zero, one, three or five minutes depending on the day, based on the performance and motivation of previous days recall sessions. Sound cue was not interrupted when the animal poked the correct port, to avoid the feedback signal that could have interfered with the memory recall. The correct port was the same than in the prior learning session, performed 2 hrs earlier on that same day. After the period with now water, the recall sessions became identical to the learning session, reinforcing the memory of that port. The recall session last 10min beyond the acclimatization period.

### Behavioral paradigm - Pre-training

this period took between 20 and 28 sessions (depending on the animal), where animals were run for one or two sessions a day. In general, after two consecutive days with performance higher than 90%, animals were passed to the next stage. The same criteria was used for every stage of the pre-training sessions. On the first day, there was a cue light on the correct port during the tone in all trials. Only on the first day there was a small water drop hanging from the water spout on the correct port once the sound cue started. Extra water was obtained when the animal poked the correct port during the tone (Fig. 1b). The sound cue lasted for up 220 secs. After approximately two days animals went above 90% performance and they were passed to a new stage. This new stage had light on the correct port, just for the first two trials, and lasted between 2-3 days. Then, the sound cue was reduced to 20 seconds in each trial. At this stage they reached criteria after 5-10 sessions. On the next stage there was not light on the first two trials indicating the correct port. In general the criteria was reached after a 5-6 sessions, where we included for this stage a minimum of 30 correct trials per session. The next stage was the last where learning session started: sound cue lasted up to 6 secs. Once they reached criteria and had more than 30 trials correct on learning sessions, animals started performing the recall session 2 hrs after the learning session on the same day.

### Analysis of Behavioral data

the output data for each session were the full video of the session recorded from a camera located above the maze, the trajectory of the animal during the entire session, the locations and times of the trigger zones, and the poke times for every port. These data were analyzed offline to characterize behavioral performance and to correlate behavioral variables with calcium imaging activity. A poke was defined as an entry into a port. We did not consider consecutive pokes in the same port unless the animal had moved 5cm away from that port. The time animals were inside one port were also saved to represent the persistency of each poke (shown as small dots in Figs. 2a and 4a), but they were not included in our analyses. Two out of twenty one animals used for the learning sessions analysis were not included in the recall sessions analysis, because their average performance across learning sessions did not exceeded 50% (chance level was 12.5%). We only included recall sessions in the analysis with a minimum of three completed trials, established as a condition on task engagement.

We quantified the accuracy of the memory recall using the memory index (MI) which was computed in three conditions: (1) 2-hours MI for single sessions and single animals (Fig. 4f-h); (2) 2-hours MI of single animals averaged across all recall sessions (Fig. 4i and Supplementary Fig. 2); (3) MI of previous days averaged across sessions and animals (Fig. 4j-l). To compute the MI of single sessions, we first computed a poke histogram of the session representing the total number of pokes *N* in each port. The port index *i* = −3, −2, −1, 0, 1, 2, 3 and 4 labeled (i.e. aligned) the different ports with respect to the correct port i=0 chosen that day (Fig. 4f). We then normalized each histogram *n_i_ = N_i_ /N*, where *N* was the total poke number in that session *N* = *N_1_* + *N_2_* + …*N_8_*. Finally, we computed the *poke vector V* that measured the net excess of poking probability (vector length) towards one spatial direction (vector angle) (see arrow in Fig. 4f-g). The vector was defined as the vector sum of the eight subvector components of length *n_i_* and angle *θ_i_*, each of them representing the tendency to poke the corresponding *i*-th port. The angles of the ports were *θ_0_* = 0, *θ_1_* = *π*/4, *θ_2_* = *π*/2, *θ_3_* = 3*π*/4, *θ_4_* = *π*, *θ_−3_* = 5*π*/4, *θ_−2_* = 3*π*/2 and *θ_−3_* = 7*π*/4. Thus, the x and y components of *V* were precisely defined as:

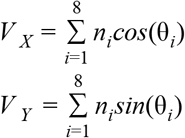

The memory index was equal to the x component of the pokes vector, MI = *V_x_*. (see red segment in Fig. 4f-g). To compute the average MI across recall sessions in a single animal, we first created a pooled poke histogram by summing the poke histogram of each session (always aligned at the correct port; Fig. 4i), then normalized the pooled histogram and then obtained a session-averaged MI (red segment in Fig. 4i), just as we did from the single session poke histograms. To obtain the MI of the rewarded ports learned on a previous day (e.g. day lag −1, −2, …-8), we realigned each single session histogram with respect to the port that had been correct on that previous day. Sessions from the same animal were then pooled, normalized and averaged across animals (Fig. 4j-k). MIs for the memory of previous days (day lag −1, −2, …-8) were then obtained from this histograms as described above (Fig. 4l).

To assess the statistical significance of the poke histograms and MIs we generated surrogate data sets representing poking activity compatible with the null hypothesis, and obtained 99% confidence bands. For single sessions, we generated M=1000 poking surrogate sets each generated from a uniform distribution of pokes across the eight ports (i.e. there were no port preferences) and having the exact same total number of pokes *N* as the data. For each surrogate set, we obtained the poke histogram and the MI as done with the original data. We then obtained a *P* < 0.01 pointwise significance bound for the poke histogram (magenta dashed line in Fig. 4f-g) and for the MI (magenta dashed line in Fig. 4h). For the session-averaged histograms, we adopted a more conservative null hypothesis: surrogate data sets were obtained by shuffling the port indices of the single session histograms, then pooled over sessions, normalized and converted into MIs. This shuffled data represents a null hypothesis in which the poking distribution deviates from a uniform distribution similarly than the original data (i.e. certain ports were preferred over others in each session), but these preference deviates show no relation with whether the port was correct or not. Finally, assessing significance of memories from previous days required to generate surrogates from a generative model representing the null hypothesis. The reason was that the location of the correct port was not independently drawn in each day and for each animal. Instead, we minimized the probability that (1) two animals running on the same box had the same correct port on the same day, and (2) that each animal had the same correct port on consecutive days. The second condition introduced serial correlations in the sequence of correct ports experienced by each animal across days. In particular, the probability that today’s correct port had been also correct on previous days was lower than chance (see inset Fig. 4l). These correlation caused that, an excess of probability to poke today’s correct port, would automatically cause an artifactual lack of probability to poke on the correct port from previous days. To get around this problem, we modeled the poking behavior of an agent who only had memory about the correct port of the same day (i.e. only 2-hours memory) with an accuracy matching the MI of each single session. In particular, the distribution of pokes of the model was uniform in all ports except the correct port in which there was the precise excess (or lack) of probability to match the MI of that day. We used this model to generate M=1000 surrogate sets for each session and each animal, using the same number of pokes and following the exact same correct port sequence as in the experiment. From the M surrogates we obtained averaged histograms for the memories of previous days. These histograms showed a deep in the center corresponding to the lack of poking probability caused by the spurious interaction of the 2-hours memory across days (see magenta dashed lines in the lower insets of Fig. 4j-k). Because of this, the MIs for previous memories obtained from these surrogates were negative for the immediately preceding days and then tended to zero as the session lag increased (gray line in Fig. 4l shows the surrogate mean MI). We finally obtained the significance bound (*P* < 0.01) from this surrogate MIs and used it to assess the significance of the original data MI (comparison between back dots and the magenta dashed line in Fig. 4l). We checked that this significant bound did not change if the null hypothesis was model using a Von Mises distribution for pokes.

### Surgeries for implantation of mini-endoscopes for calcium recordings

animals received food and water ad libitum for 2-3 days prior to the surgery. All animal procedures were approved and executed in accordance with institutional guidelines (Generalitat de Catalunya: Autorització de projecte d’experimentació N°=9997), and the Comitè Ètic d’experimentació Animal (CEEA) from University of Barcelona, N^o^=121/18.

### Viral injection

we performed surgeries when mice were between 16-18 weeks of age, once they were trained on the task and their average performance across days reached 80%. We label excitatory neurons by injecting an adeno-associated virus (AAV, serotype 2.5) driving expression of GCaMP6m via the CaMKIIα promoter. We anesthetized mice with isoflurane (induction, 5%; maintenance, 1–2%) in 95% O2, and then fixed in a stereotactic frame (Kopf Instruments). We stabilize the body temperature at 37°C using a temperature controller and a heating pad. We inject 600 nL of the AAV (injection coordinates relative to bregma in three locations: medio-lateral (ML)=1.8., anterior-posterior (AP)=−1.5, dorso-ventral (DV)=−1.6; ML=1.4., AP=−2.2, DV=−1.55; ML=2.1., AP=−2.9, DV=1.8, from bregma) via a borosilicate glass pipette with a 50-μm-diameter tip using short pressure pulses applied with a picospritzer (Parker).

### Mini-endoscope implantation

30 days after AAV injection we performed a second surgery to implant what we call the mini-endoscope, which is a stainless steel guide tube (1.2 mm diameter) with a custom glass cover slip glued to one end (0.13 mm thick cover glass, Paul Marienfeld GmbH), which holds the GRIN lens to focus then the cells on the camera of the mini-microscope. To ensure a stable attachment of the implant, once the cranium had dried we inserted 6 small screws (18-8 S/S, Small parts) on top of the cerebellum, olfactory bulbs and somatosensory sensory cortex making the largest cross possible to increase torque resistance of the implant. We perforated the skull with dental milling bit of 0.7mm diameter, and we screw-in the screws for approximately 1/2 millimeter to avoid piercing the dura. We then inserted the mini-endoscope with the position and the angle to cover the more extended flat area of the dorsal part of hippocampal CA1 region (relative to bregma ML=2.1(+77° on the coronal plane), AP=−2.2, DV=−1.1(from dura)). To perform the implantation, we first made a round craniotomy centered on the injection coordinates using trephine drill (1.6 mm in diameter). To prevent increased intracranial pressure due to the insertion of the implant, we aspirated all brain tissue inside the cilindric volume that the mini-endoscope occupies, taking out up to the second set of fibers crossing over the CA1 are, coming from the entorhinal cortex. We recognized each set of fibers by identifying their orientation that are ~60 degrees from the previous layer. Next, we lowered the mini-endoscope with the manipulator of the Kopf Table fixing the mini-endoscope to the skull using ultraviolet-light curable glue (Loctite 4305). We then applied Metabond (Parkell) around both screws, the implant and the surrounding cranium. Lastly, we applied dental acrylic cement (Coltene, Whaledent) on top of the Metabond, for the joint purpose of attaching a metal head bar to the cranium and to further stabilize the implant. Mice recovered after 3 days, but we wait up to 5-7 weeks for the tissue to return to its place after movement due to neuro-inflammatory processes. At this point we checked the level of GCaMP6m expression by locating the GRIN lens within the endoscope, we head fixed the mouse and we approach the mini-microscope to adjust focus to see neurons activate while the animal was running on a wheel. If expression was sufficiently bright, we then mounted the miniature microscope’s base-plate (nVista HD, Inscopix Inc.) utilizing acrylic cement and the ultraviolet-light curable glue (Loctite 4305).

### Calcium recordings during behavior

we brought in a second computer to control and store the frames captured by the nVista mini-microscope. We used the TTL output from the DAQ board to the arduino, to record the time-stamps of each frame captured by the mini-microscope. We then off-line combined the calcium activity with the animal behavior. We downsampled the raw movies from the miniscope prior to the processing due to computer memory constraints. We used the NoRMCorre piecewise linear registration algorithm (Pnevmatikakis et al 2017) to minimize the frame-to-frame displacements caused by the animal’s movement. Next, we took the dfof of the movies by subtracting and dividing each pixel activity at a given frame by its mean activity across the recording session. We then applied the CELLMAX extraction algorithm (Kitch L.J 2015), which models the way the movies arise from the underlying calcium signals, and finds the most likely set of neurons in the movie by doing maximum likelihood on this probabilistic generative model. Applying this algorithm on a temporally downsampled version of the dfof movies, we obtained 600 to 1000 mask candidates for the neurons in a given session. These candidates were inspected in a semi-automated manner for calcium-like dynamics and neuron-like shapes, resulting in 300-500 simultaneous neurons per session.

## Supplementary Figures

**Supplementary Figure 1:**
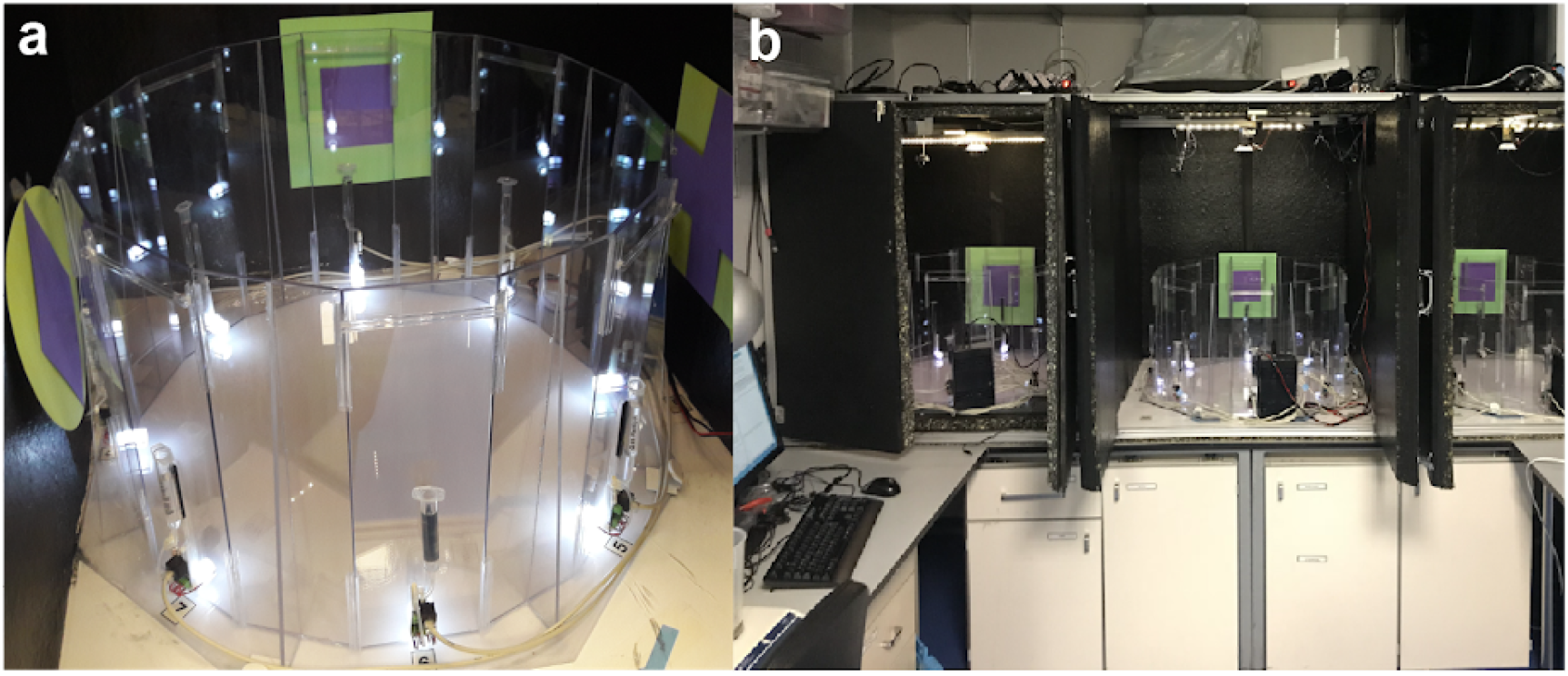
Behavioral boxes for freely moving navigation tasks. **a)** Picture of the task maze showing the transparent acrylic walls, the water ports, and the visual cards located on the inner walls of the isolation box. **b)** picture of three sound-isolating boxes placed next to each other and controlled by the computer to minimize human intervention. With these three boxes, we were able to run 18 animals per day, yielding a total of around 1400 trials per day (including learning and recall sessions).

**Supplementary Figure 2.**
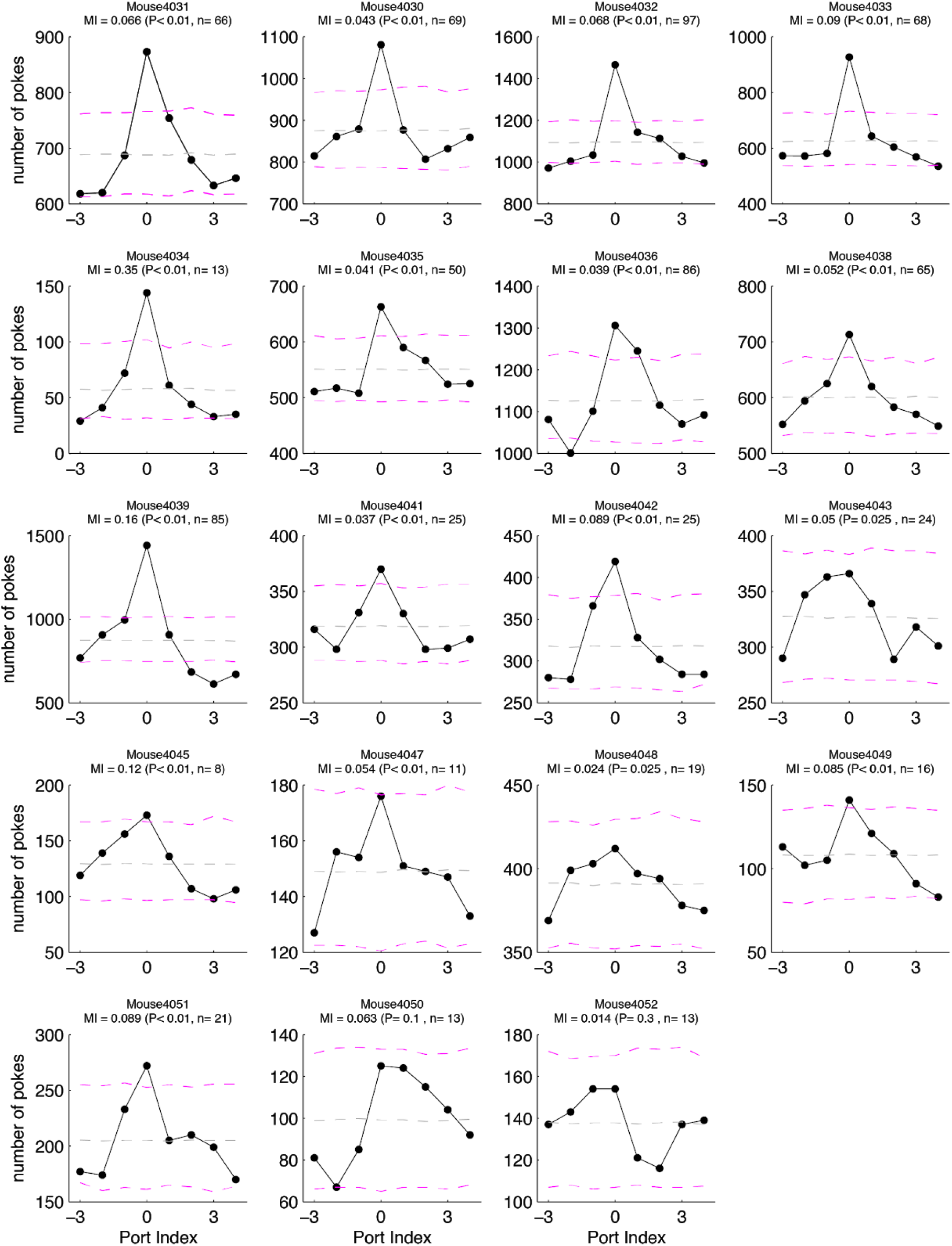
Poke histograms during recall sessions for individual animals. Poke histograms of each animal (different panels) averaged across all recall sessions (see *n* above each panel; pokes were taken from both on and off sound cue periods). Correct port index was zero. Magenta dashed lines represent a 1%-99% confidence interval obtained from shuffled surrogate data separately generated for each animal. Counts above or below this band are significantly different than chance. Gray dashed line is the mean value of the shuffled data. Memory indices (MI) obtained from each of these histograms, are indicated above each panel, together with their degree of significance.

